# Cellular consequences of non-ablative radiotherapy, a novel approach to ventricular tachycardias

**DOI:** 10.64898/2025.12.12.693887

**Authors:** Claudia Maniezzi, Stefano Busti, Angela S. Maione, Elena Sommariva, Chiara Florindi, Gaetano M. De Ferrari, Francesco Lodola, Antonio Zaza

## Abstract

**Background:** Radiotherapy (RT) with a single focused application of ionizing radiation (STAR) has been suggested as a non-invasive alternative to radiofrequency in ablating ventricular tachycardia (VT). Emerging data reveal that STAR may suppress VT without substrate destruction, by enhancing impulse conduction instead, through increased expression of NaV1.5 channels and connexin 43.

**Aims:** To investigate electrophysiology and intracellular Ca^2+^ dynamics in cardiomyocytes (CMs) from mice subjected to *in-vivo* RT. The emerging data led us to evaluate biochemical changes potentially linking electrophysiological response to ionizing irradiation.

**Methods:** CMs isolated 2 weeks after RT with low-dose (15 Gy) or high-dose (25 Gy) were compared to those of sham-treated mice (CTRL). We evaluated: i) I_NaT_ and I_Nasus_ properties; ii) AP parameters, including the prevalence of Early After-Depolarizations (EADs); iii) intracellular Ca^2+^ dynamics; iv) CaMKII phosphorylation and v) ROS content.

**Results:** 25 Gy RT i) increased I_NaT_ and, to a larger extent, I_Nasus_ (increased I_Nasus_/I_NaT_ ratio); ii) increased AP amplitude, +dV/dt_max_ and duration (APD) and facilitated EADs; iii) depressed intracellular Ca^2+^ dynamics. 15 Gy RT had similar but smaller effects (dose-dependency). 25 Gy RT reduced CaMKII phosphorylation but increased cell ROS content, thus providing a mechanism for I_NaL_ enhancement.

**Conclusions:** The results support the view that STAR may supress VT by increasing conduction velocity, with APD prolongation providing an additional mechanism. On the other hand, I_Nasus_ enhancement (likely by ROS) and Ca^2+^ handling depression may impair electrical stability and contractility in the irradiated region.

## Introduction

Ventricular tachycardias (VTs) are a frequent complication of advanced structural heart disease because of a complex combination of anatomical and functional predisposing factors. Besides being a life-threatening condition, VTs may be symptomatic and, due to the risk of inducing syncope, a strong limitation to patients’ lifestyle.

VTs treatment often requires anatomical ablation of the arrhythmogenic substrate, mostly a reentry circuit, currently performed by local application of thermal energy (catheter ablation, CA). Albeit useful, the procedure is invasive, major complications occur in up to 5-6% of procedures ^1^ and VT recurrence rate is between 27% and 55% depending on the substrate ^2^. Furthermore, the scar generated by the procedure might provide a new arrhythmogenic substrate.

External application of ionizing radiation to ablate the VT substrate has been proposed in 2013 as a non-invasive alternative to CA ^3^ and has been evaluated in small clinical studies with positive and enduring results ^4-6^. This approach, named Stereotactic Arrhythmia Radioablation (STAR), is based on a single delivery of high-dose (25 Gy) radiotherapy (RT) to a relatively small volume (mean 25 cc) of myocardium by a focused beam.

As conceived, STAR is still “ablative”, i.e. meant to replace the tissue supporting the arrhythmia with an unexcitable fibrotic scar. Nonetheless, VT suppression was observed at times after STAR delivery hardly compatible with scar formation ^4^, thus suggesting a functional mechanism instead. A recent article by Zhang et al ^7^ indeed reported the absence of fibrosis in the irradiated tissue from human hearts in which VT was successfully treated by 25 Gy STAR. When applied to mice, the irradiation increased myocardial conduction velocity, enhanced the expression of Na^+^ channel proteins (NaV1.5) and of the gap-junctional protein Cx43. This finding opens the theoretically plausible possibility to treat VT by boosting impulse propagation, as opposed to discontinuing it by tissue damage. Since this approach aims to preserve viability of irradiated myocytes, its effects on myocyte electrophysiology, ionic homeostasis and biology deserve investigation.

Enhanced expression of NaV1.5 channel protein expectedly increases the Na^+^ current (I_Na_); however, evidence on this aspect is missing. Moreover, whether the transient (I_NaT_) and sustained (I_Nasus_) components of I_Na_ are equally affected is of interest. Indeed, whereas I_NaT_ upregulation is crucial for conduction improvement, I_Nasus_ enhancement may affect membrane potential and intracellular Ca^2+^ homeostasis, with undesirable effects on electrical stability, tissue remodelling and energetic competence^8^.

In the present study we investigate the effects of cardiac RT at two doses (25 and 15 Gy) in murine cardiomyocytes with the aim to verify the results of Zhang et al by direct action potential and I_Na_ measurements (not available in Zhang et al ^7^) and to evaluate RT effect on intracellular Ca^2+^ dynamics. The 25 Gy dose is the one used in the previous studies providing the background for the present one, 15 Gy irradiation was also tested to establish if the molecular and functional changes responsible of VT suppression could be achieved with a lower dose.

## METHODS

### Irradiation protocol

Whole heart X-ray irradiation was performed at the National Institute of Nuclear Physics (INFN-Catania) within the Reasearch Project PRIN 2022 “*Non-ablative radiotherapy in cardiac arrhythmias: a new paradigm beyond scar induction*” funded by The Italian Ministry Of University and Research, Protocol # 2022L7H8B4. Approval of the irradiation procedure by local and national Ethical Committees was obtained during the Project application and is implicit in its approval.

Lots of irradiated mice plus matched controls were shipped to our laboratory within one week from irradiation at a pace compatible with execution of all experiments at the established timeline.

### Electrophysiology

Mouse myocytes (CMs) were studied 2 weeks after whole-heart irradiation (25 Gy and 15 Gy) and compared to non-irradiated littermates (controls, CTRL). Ventricular myocytes were enzymatically isolated and studied by patch-clamp after Ca^2+^ adaptation. Cells were superfused with Tyrode’s solution (36.5 °C) containing (mM): NaCl 154, KCl 4, CaCl_2_ 2, MgCl_2_ 1, HEPES 5, Glucose 5.5, pH=7.35 titrated with NaOH. The pipettes (tip resistance of 2-3 MΩ) were filled with a solution containing (mM): K-Aspartate 110, KCl 23, MgCl_2_ 3, CaCl_2_ 0.04, EGTA KOH 0.1, HEPES KOH 5, Na-CP 5, Na-ATP 5, Na-GTP 0.4, pH=7.3 with KOH.

Evaluation of the I_Nasus_/I_NaT_ ratio required to measure both I_Na_ components within each myocyte. Physiological extracellular concentrations and temperature may reduce the accuracy of I_NaT_ measurements (potential V_m_ bias due to series resistance), nonetheless they are required for adequate resolution of I_Nasus_. Since characterization of I_NaT_ V-dependence was not an aim of the study, we prioritized accurate I_Nasus_ quantification. To this end, both I-clamp and V-clamp measurements were performed under superfusion with normal Tyrode’s solution (above) at 36.5 °C.

#### Action potentials (AP) recordings

APs were recorded (I-clamp) during steady stimulation at 1 Hz (2 ms pulses at twice the threshold through the pipette). The following AP parameters were measured: diastolic potential (E_diast_), action potential amplitude (APA), maximal upstroke velocity (+dV/dt_MAX_), action potential duration at 90% repolarization (APD_90_). The prevalence of early afterdepolarizations (EADs), arrhythmogenic repolarization abnormalities typically associated with I_Nasus_ enhancement, was measured as the % of cells displaying them. Finally, to verify I_Nasus_ involvement in AP changes, AP response to selective I_Nasus_ blockade (1 µM tetrodotoxin, TTX) was assessed.

#### Na^+^ current (I_Na_) recording

A step-ramp voltage protocol was used to quantify transient (I_NaT_) and sustained (I_Nasus_) components respectively. I_NaT_ was recorded during a single step from -90 to 0 mV. I_Nasus_, contributed by both “window” and “late” (I_NaL_) components, was recorded during a V ramp of velocity mimicking repolarization (from +40 to -100 mV, 56 mV/s) and dissected by subtraction as the current sensitive to 30 µM TTX; this protocol allowed to construct quasi-steady state I/V curves of the current; I_Nasus_ reversal potential (E_rev_) was obtained by linear fitting of the ascending portion of the curves. Current density was obtained by normalization to cell capacitance.

### Intracellular Ca^2+^ dynamics

Intracellular Ca^2+^ activity was measured by dynamic epifluorescence under field-stimulation at constant rate (1 Hz) and physiological temperature. To this end, myocytes were loaded with the Ca^2+^-sensitive (Fluo-4 AM, 10 μM) fluorophore. Fluorescence (F) was normalized to the value measured after prolonged quiescence (F0). The following parameters of Ca^2+^ transients (CaT) were quantified: i) CaT amplitude, ii) CaT time to peak (t_peak_), iii) maximal rate of CaT rise (dCa/dt_MAX_), iv) CaT decay kinetics (τ decay), and v) diastolic Ca^2+^ (CaD). Rate-dependency of CaT properties was tested by incrementing the pacing rate from 1 to 5 Hz.

### Biochemical assays

1. CaMKII activation was assessed by western blot of homogenates of cell pellets from ventricles, evaluating the ratio between phosphorylated CaMKII and unphosphorylated one. Densitometric analysis was performed by ImageJ software (see Supplement for detail).
2. Intracellular ROS concentration was measured in single cells after incubation with the fluorescent probe 2′7′-dichclorofluorescin diacetate (DCFDA, 10 μM). Brightfield and fluorescence images were acquired with a Leica THUNDER LIVE imaging system (LAS X software) using a 20X objective in a controlled environment (37°C, 5% CO2). Cell segmentation (identification of single cells) was performed on brightfield images by using a custom version of the Cellpose cyto3 model ^9^ specifically trained to recognize cardiomyocytes (efficiency > 95%). Live/dead cells were distinguished (and excluded from ROS quantification) based on their different morphology by using the ilastik machine-learning classifier ^10^. Fluorescence intensities were extracted for each single cell by using Fiji/ImageJ.

### Experimental design

A “unpaired sample” design was adopted for comparing the effects of irradiation vs non-irradiation (control) and 15 Gy vs 25 Gy irradiation.

### Statistical analysis

Statistical analysis was carried out with GraphPad Prism 8 and R softwares.

Normality of distribution was assessed using D’Agostino-Pearson’s normality test. One-way ANOVA (with the respective post-hoc corrections) was used for multiple comparisons of continuous variable. In the case of repeated measurements (e.g., rate-dependency) a mixed-effects ANOVA model containing “Treatment” (CTRL vs 15 Gy vs 25 Gy) and “Variable” (pacing rate) factors was used. Significance of “Treatment × Variable interaction” (i.e., difference between Treatments in their response to the Variable) was first tested; in its absence, significance of difference between Treatments at all Variable values was tested. Fisher exact test was applied to contingency tables for multiple comparison of categorical variables (EADs prevalence); the Benjamini-Hochberg correction was applied for post-hoc comparisons between individual frequencies.

In figures, data are presented as mean ± SEM. For each experiment, the number of preparations or cells (n) and the number of animals from which they were obtained (N) are indicated in the respective figure legend.

## RESULTS

### 1.1 Effects of RT on transient (I_NaT_) and sustained (I_Nasus_) components of I_Na_

The effect of RT on I_NaT_ and I_Nasus_ is illustrated in **Fig. 1**. 25 Gy RT increased both I_NaT_ (+65%, p<0.05), **Fig. 1A**) and I_Nasus_ (+242%, p<0.0001, **Fig. 1B** and **1C**), as expected from an increased expression of Na^+^ channels. Beside increasing maximal I_Nasus_ density, irradiation (at both 15 and 25 Gy) shifted by about +10 mV the potential at which maximal current was achieved, thus increasing current amplitude preferentially at potentials positive to -30 mV (**Fig 1B**). The I_Na_ “window” encopasses potentials negative to -30 mV ^11 12^; therefore, the finding suggests that the increase in I_Nasus_ was contributed by its I_NaL_ component. Notably, the % increments in I_NaT_ and I_Nasus_ differed, thus resulting in a significant increase in the I_Nasus_/I_NaT_ ratio (+159%, p<0.001 **Fig. 1D**). E_rev_ was +45.6±1.8 mV in control and was unchanged by irradiation (**Fig. 1E**), thus confirming that I_Nasus_ was carried by Na^+^ in all the experimental conditions.

**Figure 1.**
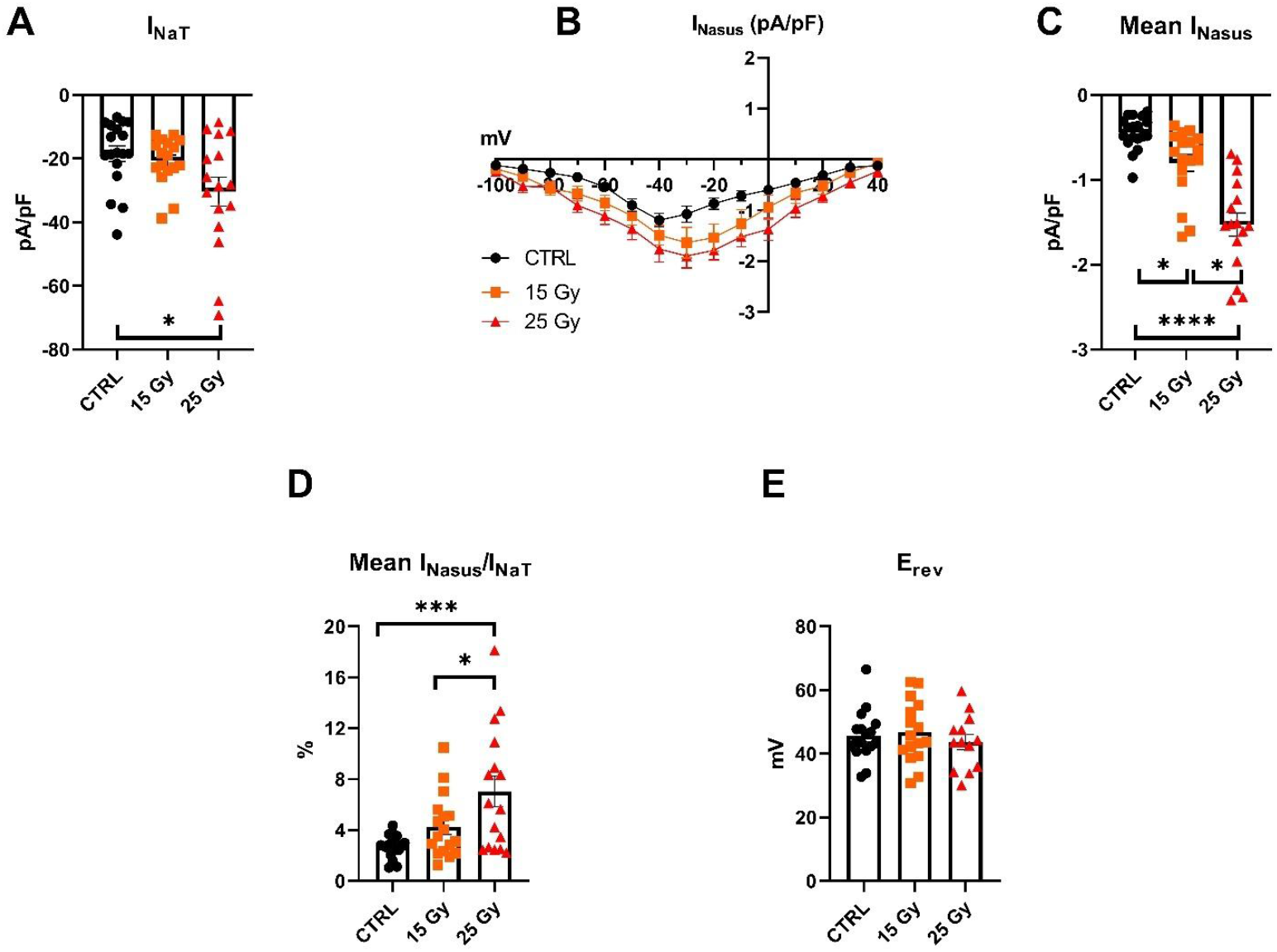
Effect of high dose (25 Gy) and low dose (15 Gy) RT on transient (I_NaT_) and late (I_Nasus_) component of I_Na_. (**A**) I_NaT_. (**B**) I_Nasus_ current-to-voltage relationship. (**C**) Mean I_Nasus_. (**D**) Mean I_Nasus_/I_NaT_ ratio. (**E**) Reversal potential (E_rev_). CTRL: n=18, N=3; 15 Gy: n=17, N=5; 25 Gy: n=16, N=2. Data are expressed as mean ± SEM. One-way ANOVA, * p<0.05, ** p<0.01, *** p<0.001, **** p<0.0001.

With 15 Gy, the changes were smaller (dose-dependency) and achieved statistical significance for I_Nasus_ only (+79%, p<0.05, **Fig. 1C**) thus tending to increase the I_Nasus_/I_NaT_ ratio (**Fig. 1D**).

Overall, the results show an I_NaT_ increment compatible with the effect of irradiation on myocardial conduction velocity ^7^, but also a preferential increase in I_NaL_, of potentially pathogenetic relevance.

### 1.2 Effect of RT on the action potentials (AP)

AP parameters were measured at steady state during 1 Hz stimulation (**Fig. 2**). 25 Gy RT enhanced APA (+12%, p<0.05, **Fig. 2B**) and +dV/dt_MAX_, (+38% p<0.05, **Fig. 2C**), consistent with the increment in I_NaT._ It also significantly prolonged APD_90_ (+110%, p<0.01, **Fig. 2D**), as expected from the increment in I_Nasus_; E_diast_ remained unchanged (**Fig. 2E**). 15 Gy RT failed to modulate APA (**Fig. 2B**) and +dV/dt_MAX_ (**Fig. 2C**) but significantly prolonged APD_90_ (+69%, p<0.05, **Fig. 2D**). EADs prevalence was 12% in control CMs, 29% in 15 Gy-irradiated CMs and 37% in 25 Gy-treated CMs (**Fig. 2F**). Despite its dose-dependency, the increment in EADs prevalence failed to achieve statistical significance (p>0.05 at Fisher test). EADs occurred at a membrane potential around -40 mV (plateau phase, **Fig. 2A**) where I_Nasus_ was maximal (see **Fig. 1B**), thus suggesting its direct contribution to EADs.

**Figure 2.**
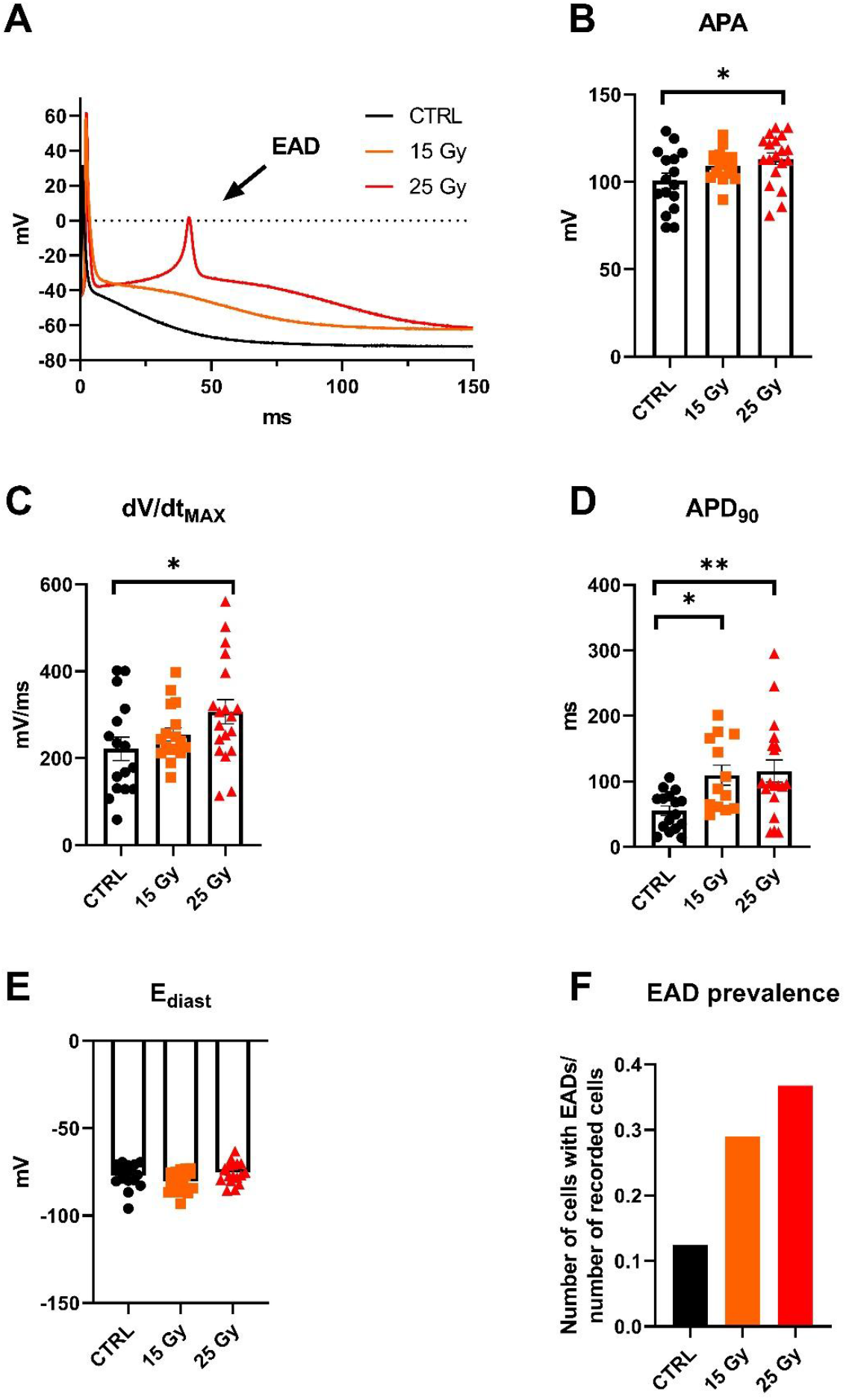
Effect of 15 and 25 Gy RT on APs. (**A**) Representative traces of AP recorded during stimulation (through the pipette) at 1 Hz. Early Afterdepolarization (EAD) indicated by arrow. (**B**) AP amplitude (APA). (**C**) Maximal upstroke velocity (dV/dt_MAX_). (**D**) AP duration at 90% repolarization (APD_90_). (**E**) Diastolic potential (E_diast_). (**F**) EAD prevalence. CTRL: n=16, N=3; 15 Gy: n=17, N=3; 25 Gy: n=19, N=3. Data are expressed as mean ± SEM. One-way ANOVA, * p<0.05, ** p<0.01, *** p<0.001, **** p<0.0001.

To test whether I_Nasus_ enhancement contributed to repolarization slowing, we investigated the impact of selective I_Nasus_ blockade (by 1 µM TTX) on APD_90_ (**Fig. 3**). TTX shortened APD_90_ more in RT CMs (-11% in 15 Gy and -25% in 25 Gy) than in CTRL ones (-6%), thus confirming the role of I_Nasus_ modulation on APD_90_ prolongation by RT.

**Figure 3.**
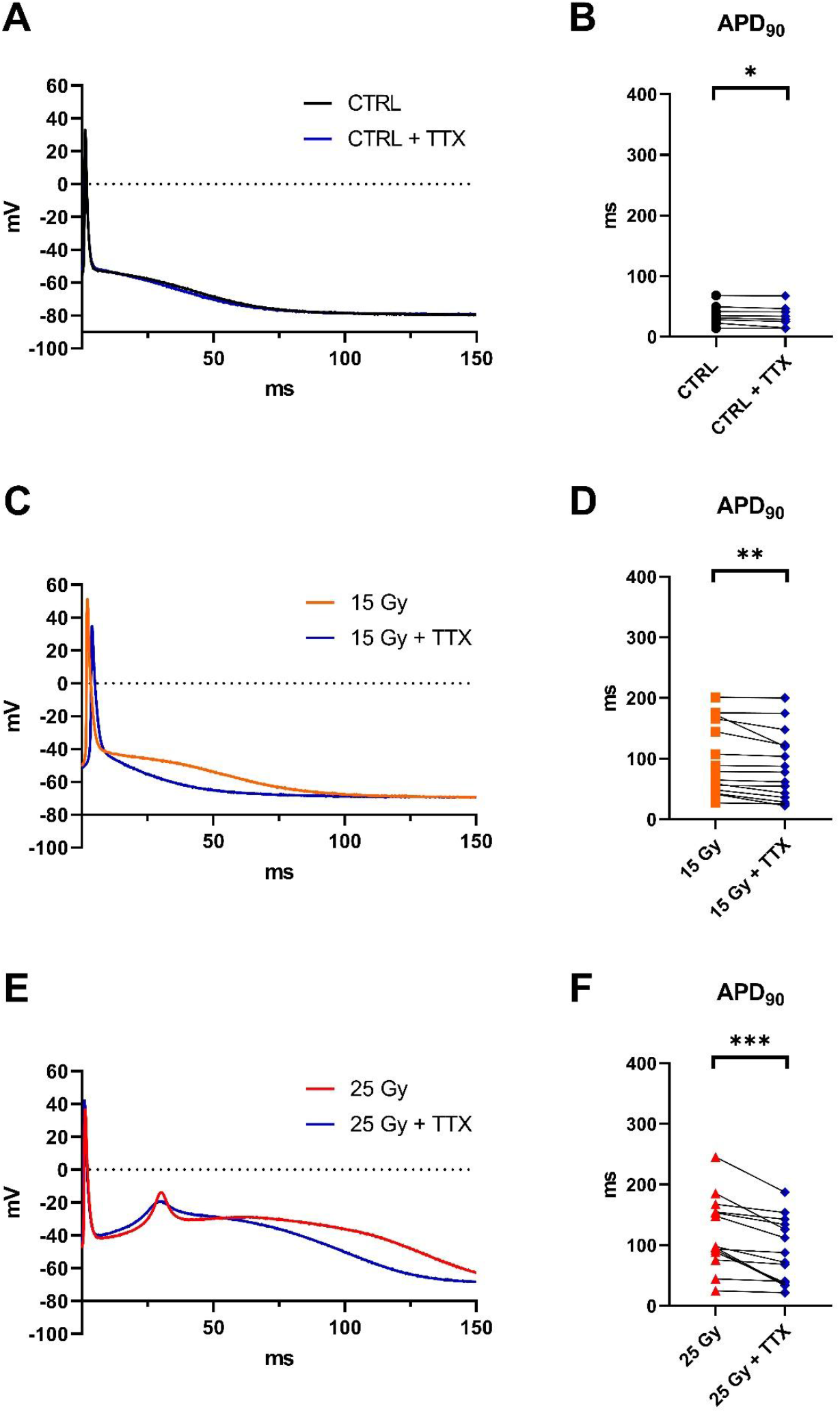
Effect of selective I_Nasus_ blockade (1 µM TTX) on APD_90_. (**A, C, E**) Representative traces of AP recorded during stimulation at 1 Hz. (**B, D, F**) APD90. CTRL: n=8, N=3; 15 Gy: n=15, N=3; 25 Gy: n=14, N=3. Data are expressed as mean ± SEM. Paired Student’s t test., * p<0.05, ** p<0.01, *** p<0.001, **** p<0.0001.

Overall, irradiation modified AP parameters in a way fully compatible with the observed increments in I_NaT_ and I_Nasus_. The substantial APD prolongation may represent a further mechanism supporting VT suppression.

### 1.3 Effect of RT on intracellular Ca^2+^ dynamics

Since Na^+^ influx through I_Nasus_ affects intracellular Ca^2+^ handling, we evaluated the effect of RT on cytosolic Ca^2+^ dynamics in field-stimulated CMs.

#### 1.3.1 Steady-state stimulation (1 Hz)

CaT parameters were quantified during steady-state stimulation at 1 Hz (**Fig. 4**). In 25 Gy-irradiated CMs, CaT amplitude was significantly reduced (-56% p<0.0001, **Fig. 4C**) while CaD remained unchanged (**Fig. 4B**). CaT kinetics were generally slowed by RT, as demonstrated by the increase in t_peak_ (+10%, p<0.01, **Fig. 4D**), the reduction in dCa/dt_MAX_ (-10% p<0.05, **Fig. 4E**) and the increase in τ decay (+20%, p<0.05, **Fig. 4F**). In 15 Gy-irradiated CMs, a similar trend in CaT parameters was observed, but the changes were smaller and did not achieve statistical significance (**Fig. 4**).

**Figure 4.**
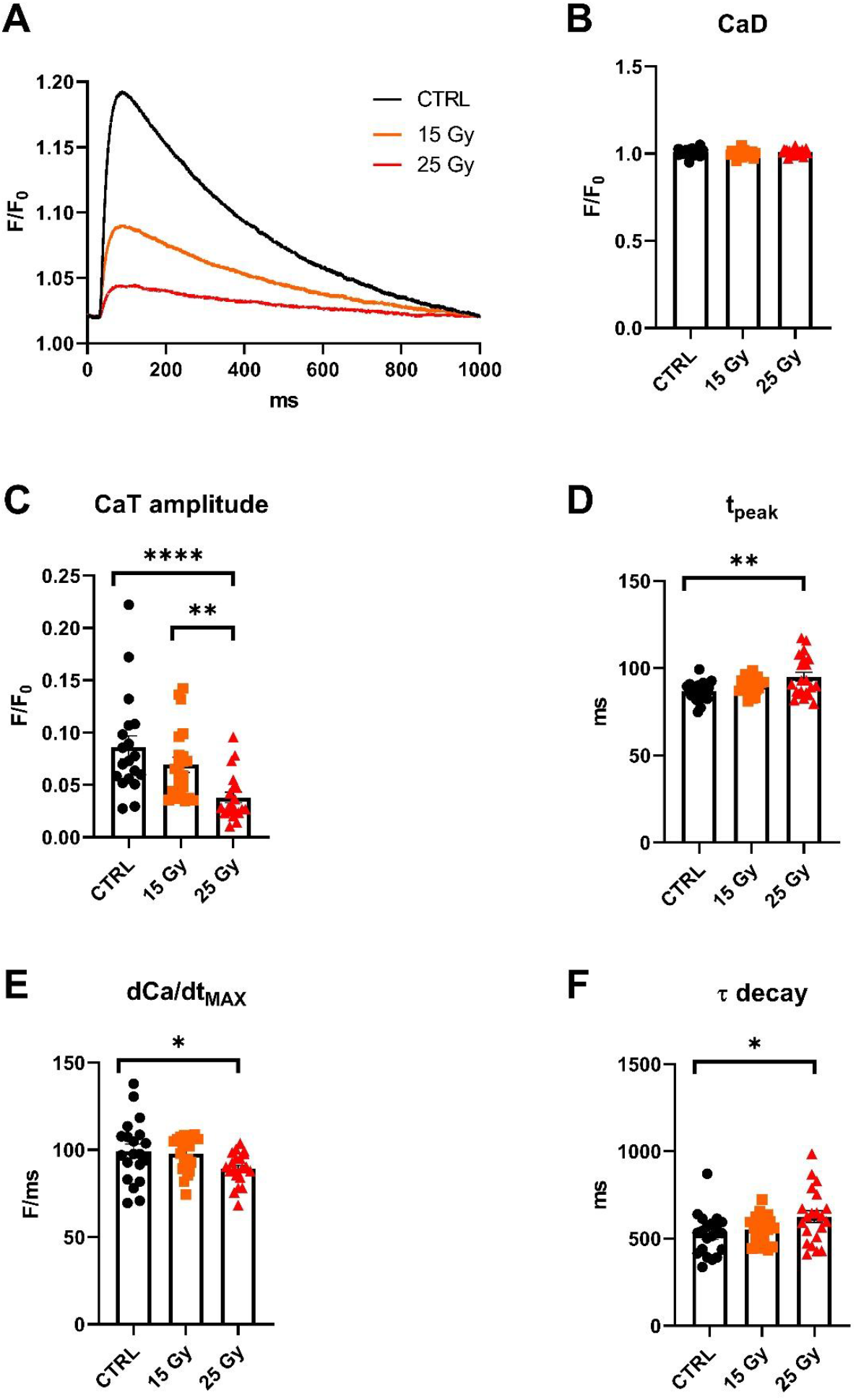
Effect of 15 and 25 Gy RT on Ca^2+^ transients (CaT) during steady-state stimulation at 1 Hz. (**A**) Representative CaT traces. (**B**) Diastolic Ca^2+^ (CaD). (**C**) CaT amplitude. (**D**) CaT time to peak (t_peak_). (**E**) Maximal rate of CaT rise (dCa/dt_MAX_). (**F**) CaT decay kinetics (τ decay). CTRL: n=20, N=2; 15 Gy: n=22, N=3; 25 Gy: n=21, N=2. Data are expressed as mean ± SEM. One-way ANOVA, * p<0.05, ** p<0.01, *** p<0.001, **** p<0.0001.

Overall, the results suggest a general downregulation of EC coupling, compatible with simultaneous impairment of various among its components.

#### 1.3.2 Rate-dependency

Rate-dependency of CaT properties was tested in a subset of cells by incrementing the pacing rate from 1 to 5 Hz.

Under control conditions, CaT parameters showed significant rate-dependency (positive or negative) (**Fig. 5**): CaD (positive, p < 0.0001), dCa/dt_MAX_ (positive, p < 0.0001), CaT amplitude (negative, p < 0.001), *t*_peak_ (negative, p < 0.0001), τ_decay_ (negative, p < 0.0001).

**Figure 5.**
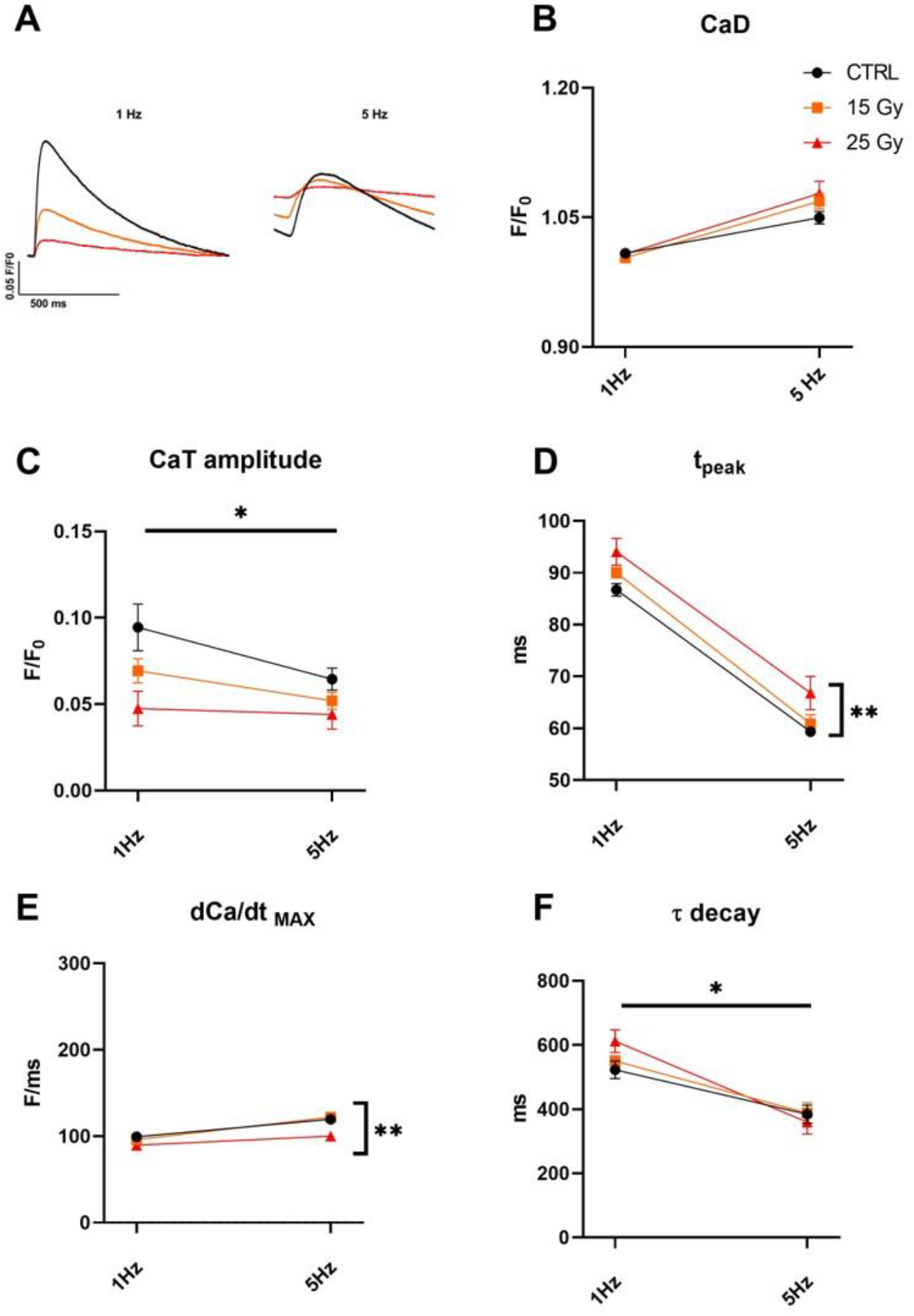
Rate dependency of CaT parameters. Rate-dependency was tested by incrementing the pacing rate from 1 to 5 Hz. (**B**) Diastolic Ca^2+^ (CaD). (**C**) CaT amplitude. (**D**) CaT time to peak (t_peak_). (**E**) Maximal rate of CaT rise (dCa/dt_MAX_). (**F**) CaT decay kinetics (τ decay). CTRL: n=20, N=2; 15 Gy: n=22, N=3; 25 Gy: n=19, N=2. Data are expressed as mean ± SEM. Mixed-effects model, Treatment X Rate.

As compared to CTRL, in 25 Gy CMs: CaT amplitude had a shallower rate-dependency (**Fig. 5C**, Treatment × Rate, p<0.05) due to preferential reduction at slower rates; τ_decay_ had a steeper rate-dependency (**Fig. 5F** Treatment × Rate, p<0.05) due to preferential increase at slower rates. As expected from I_Nasus_ enhancement, CaD accumulation at high rate tended to be more prominent in irradiated CMs (**Fig. 5B**). For t_peak_ (**Fig. 5D**) and dCa/dt_MAX_ (**Fig. 5E**), rate-dependency remained unchanged between CTRL and 25 Gy.

With 15 Gy irradiation, the same trends were observed (**Fig. 5**).

### 1.4 Mechanism for I_Nasus_ enhancement

#### 1.4.1 Evaluation of CaMKII phosphorylation

The results so far indicate that 25 Gy RT enhanced I_Nasus_, thus prolonging APD and altering the intracellular Ca^2+^ dynamics. In order to investigate the mechanisms for I_Nasus_ enhancement, we first quantified CaMKII activation by measuring its total and phosphorylated (pCaMKII) fractions by western blotting (**Fig. 6**). In RT CMs, the pCaMKII/ CaMKII ratio showed a dose-dependent trend to decrease, with a significant reduction at the 25 Gy dose, (**Fig. 6B**), which rules out CaMKII activation as a factor in I_Nasus_ enhancement.

**Figure 6.**
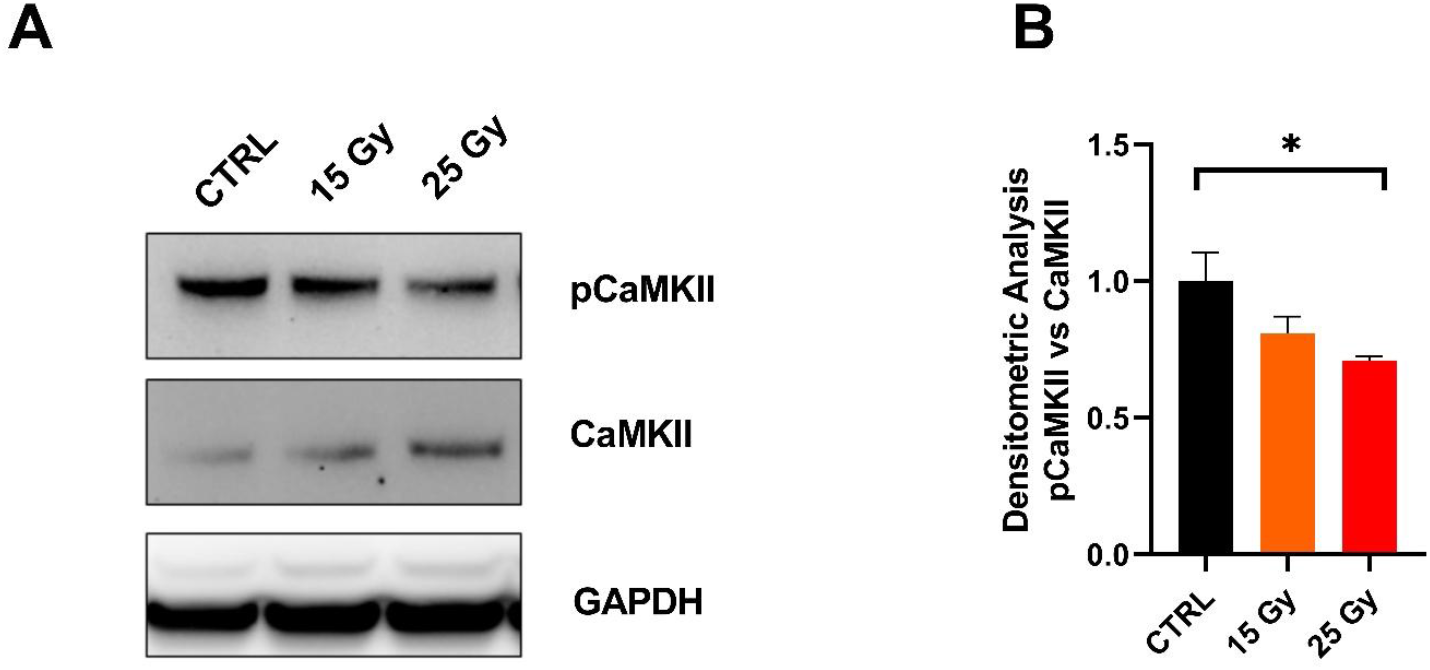
Protein analysis in cell pellet from ventricles. (**A**) Representative images of Western Blot analysis of proteins extracted from ventricular tissue. Immunostaining of the housekeeping GAPDH is shown for normalization of total CaMKII. (**B**) Densitometric analysis of the pCaMKII/CaMKII ratio. CTRL: N=3; 15 Gy, N=3; 25 Gy: N=3, Fisher p>0.05. Data expressed as mean ± SEM.

#### 1.4.2 Evaluation of cell ROS content

The intracellular content of reactive oxygen species (ROS) was estimated in quiescent CMs by measuring 2′7′-dichclorofluorescin diacetate (DCFDA) fluorescence from individual vital cells (identified by the rod-shaped contour). The analysis was limited to the 25 Gy dose, the one consistently exerting significant effects on functional parameters (above). In 25 Gy CMs, DCFDA signal was significantly stronger than in CTRL CMs (+41%, p<0.0001 **Fig. 7B**), to indicate an increase in ROS content, a likely cause of I_Nasus_ enhancement and, possibly, depression of EC coupling.

**Figure 7.**
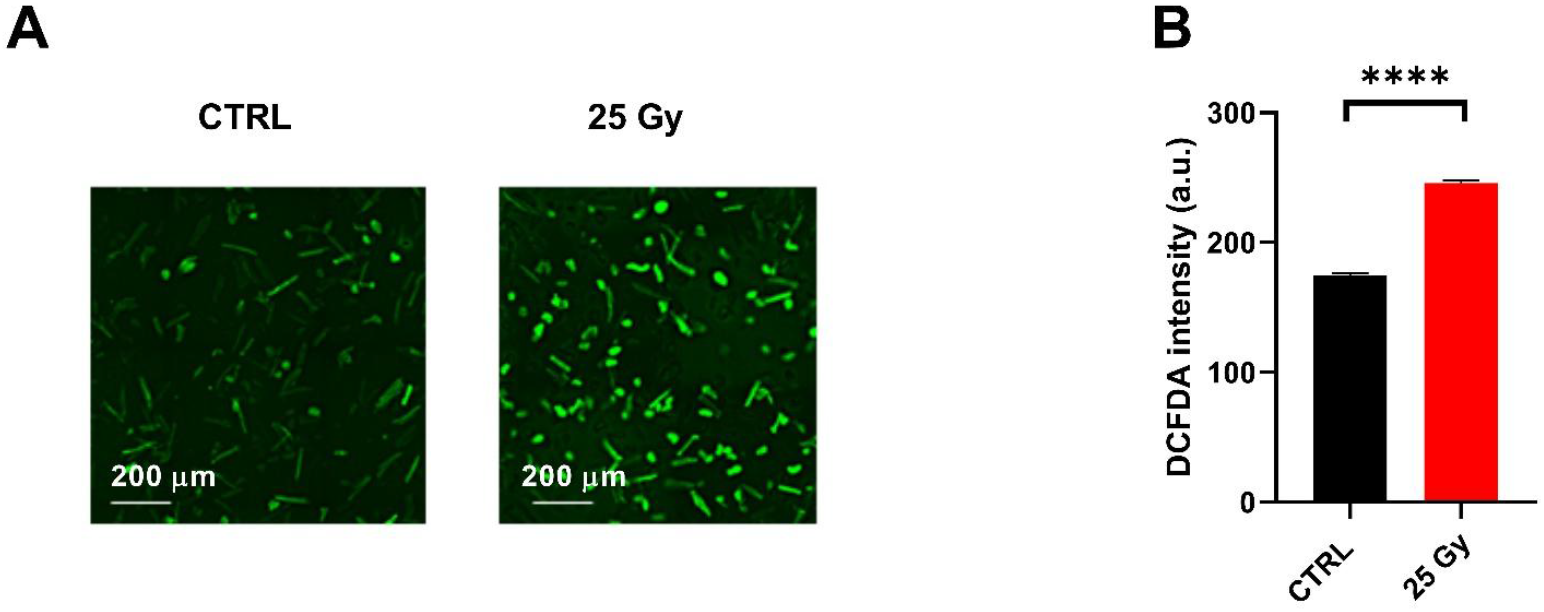
ROS quantification. (**A**) Fluorescence microscopic image (20 X) of CMs incubated with 10 µM DCFDA. (**B**) DCFDA fluorescence intensity. CTRL: N=2; 25 Gy: N=2; p<0.0001 at unpaired Student’s t test. Data expressed as mean ± SEM.

## DISCUSSION

The results of this study show, by direct I_Na_ measurement, that a single myocardial irradiation at 25 Gy may substantially increase the activity of V-gated Na^+^ channels, previously inferred from the increase in the expression of NaV1.5 protein levels ^7^. They also show that the effect concerns both the transient and sustained I_Na_ components, with a preferential effect on the latter. An increase in ROS content provides a likely mechanism for selective I_Nasus_ increment. Irradiation-induced I_Na_ changes translate into significant APD prolongation and EADs facilitation. Irradiation was also associated with significant perturbation of intracellular Ca^2+^ handling; however, its direction is only partly consistent with what expected from increased Na^+^ influx and prolonged repolarization.

### I_Na_ modulation

The observed changes in overall I_Na_ are compatible with a radiation-induced increment in membrane expression of functional NaV1.5 channels ^7^; however, the latter should increase the transient and sustained components of the current by a similar proportion. The observation of preferential I_Nasus_ enhancement suggests an additional effect of irradiation. Peroxidation of NaV1.5 channels by ROS is a well-established mechanism for enhancement of “late” I_Na_ (I_NaL_) ^13^, the “pathological” contributor to I_Nasus 8_. Two observations suggest that I_NaL_ may account for the observed I_Nasus_ increment: i) after irradiation maximal I_Nasus_ was achieved at more positive potentials, at which I_NaL_ prevails over the “window” component; ii) ROS content was indeed increased by irradiation. I_NaL_ enhancement in cardiac myocytes has also been attributed to increased expression of neuronal NaV isoforms characterized by incomplete inactivation ^14^. Since irradiation has been shown to affect channel transcription ^7^, such a mechanism would be plausible; however, the observation of a clear-cut increase in ROS points instead to post-transcriptional modulation of NaV1.5 proteins.

### AP modulation

As shown by its sensitivity to I_Na_ blockade, repolarization slowing by irradiation is attributable to I_Nasus_ enhancement. APD prolongation may contribute to the effect of irradiation in contrasting ways: i) by prolonging refractoriness it increases the wavelength of reentrant circuits; along with the increment in conduction velocity, this may contribute to reentry interruption, i.e. the desired effect of irradiation; ii) by facilitating EADs it may increase the arrhythmogenic potential of the irradiated area ^15^.

EADs facilitation by irradiation may simply result from repolarization slowing ^16^, but it might also reflect spontaneous Ca^2+^ release event promoted by ROS-induced facilitation of RyRs opening ^17^. Indeed, the observation of a perturbed Ca^2+^ handling not entirely consistent with enhanced Na^+^ influx (see below) may suggest an additional (possibly ROS-mediated) radiation effect including destabilization of the Ca^2+^ store.

### Ca^2+^ handling perturbation

By increasing Na^+^ influx, I_Nasus_ enhancement is expected to hamper Ca^2+^ extrusion, thus increasing overall intracellular Ca^2+^ content. This should lead to an increment in diastolic Ca^2+^ and, unless associated with major destabilization of the Ca^2+^ stores, to larger CaT amplitude.

While a trend to rate-dependent diastolic Ca^2+^ accumulation was indeed observed in irradiated CMs (**Fig. 5B**), CaT amplitude was conspicuously reduced in a dose-dependent fashion. This suggests that Ca^2+^ handling was perturbed by irradiation also independently of I_Nasus_ enhancement. The function of practically all the elements of the Ca^2+^ handling machinery can be affected by peroxidation ^18^. Therefore, I_Nasus_ enhancement and Ca^2+^ handling perturbation might result from the increment in cell ROS content as parallel effects, rather than being linked in a causative chain. Notably, RyRs peroxidation is a well-established mechanism of EADs facilitation ^17^.

### Increased ROS content

Redox imbalance is a known effect of ionizing radiations ^19^; therefore, the observed increment in ROS content is unsurprising. Such increment was observed 15 days after the treatment, thus qualifying as a rather sustained action, possibly accounting for irradiation effects beyond the increment in I_Na_ density.

### Dose-dependency of irradiation effects

All parameters were affected in the same direction by irradiations at 25 Gy and 15 Gy with a clear-cut dose-dependency. Therefore, failure to achieve statistical significance of the 15 Gy effects on some parameters should not be necessarily interpreted as a lack of effect, but rather as a consequence of their smaller magnitude. This view is supported by the observation that, albeit not achieving significance, the I_NaT_ increment induced by 15 Gy was associated with increments of APA and +dV/dt_max_ means, as expected from their dependency on I_NaT_ amplitude. On the other hand, 15 Gy irradiation did significantly increase I_Nasus_ and prolong APD_90_, thus suggesting higher sensitivity of these two parameters. Although the I_Nasus_ increment may be considered an untoward effect, APD90 prolongation would still support VT interruption. Overall, albeit less effective than 25 Gy, the 15 Gy dose might still exert biological effects compatible with discontinuation of reentrant circuits, but of potential proarrhythmic risk.

### Relevance to STAR therapy

As discussed above, I_NaL_ enhancement, EADs facilitation and depression of EC coupling may represent untoward effects of irradiation. The issue is to which extent these effects may dispute the therapeutic use of STAR.

The impact on overall cardiac function of the abnormalities disclosed by the present results may depend on the mass of muscle affected by the irradiation applied in the STAR procedure. Based on recent EHRA/HRS consensus data ^20^, clinically used STAR plans report median central target volumes (CTV, receiving the irradiation dose) of ∼35–50 ml and, once motion-management margins (5–10 mm) are considered, a peripheral target volumes (PTV, including the tissue receiving scattered radiation) of ∼80–90 ml. Thus, in the case of a 25 Gy dose, a CTV of approximately 40– 50 ml receives 25 Gy and an additional ∼40–50 ml of tissue surrounding the CTV is exposed to radiation doses < 25 Gy, tapering with distance.

Considering a muscle density of 1.06 g/ml a CTV of 50 ml corresponds to 53 g of muscle mass (≅ 50% of the left ventricular mass index in male), and an even larger mass is included in the PTV. The irradiated myocardium (whose excitability is enhanced) might become a source of EAD-dependent propagated activity and/or steep repolarization gradients, thereby representing a proarrhythmic risk. Such risk would depend on the affected myocardial mass, thus making beam focusing of pivotal importance.

While VT prevention (by conduction boost) may depend on I_NaT_ upregulation by transcriptional reprogramming ^7^, the proarrhythmic abnormalities may be linked to I_NaL_ enhancement instead, to which ROS may contribute significantly. If the oxidative response to irradiation were relatively short-lived, this might explain the frequent VT recurrence during the “blanking period” (6 weeks after STAR), which may subside in the subsequent months ^4^. Notably, the mechanism underlying post-procedure VT exacerbation might theoretically differ from that of the VT to be treated. I_NaL_ can be selectively blocked by suitable class I antiarrhythmic agents (e.g. mexiletine, ranolazine, amiodarone), which would remove the associated risk but, at the same time, prevent APD prolongation. Should the latter contribute significantly to discontinuation of the reentrant circuit responsible for VT, a plausible but unproven hypothesis, the corrective measure might theoretically limit STAR efficacy in VT treatment.

Currently, myocardial conduction can be improved only through gene-therapy approaches of scarce clinical applicability. Because of its effect on I_NaT_, irradiation might have an appeal as a putative treatment of conditions characterized by a diffuse conduction impairment (e.g. loss of function NaV mutations, infarct border zone etc.). However, in this case irradiation should be extended to even larger tissue volumes, potentially aggravating the side effects disclosed by the present study.

## Conclusions

The effects of a single irradiation dose of 25 Gy on I_NaT_ are consistent with the increased expression of NaV1.5 protein and account for the improvement of conduction^7^. By prolonging APD, simultaneous I_NaL_ enhancement may provide an additional mechanism for VT interruption, but it may render the (still excitable) irradiated myocardium a source of ectopic activity and weakened contraction. This may require consideration in the therapeutic application of highly focused irradiation (STAR) but might establish a contraindication to the potential use of more diffuse cardiac irradiation.

## Acknowledgement

The study was funded by PRIN 2022 (Italian Ministry of Universiy and Research) to AZ (project Coordinator GMDF) and Funds for Academic Research (FAR UNIMIB) 2022-2025 to AZ and FL. Funds for “Ricerca Corrente” of Centro Cardiologico Monzino IRCCS (Italian Ministry of Health) to ES.

## Authors’ role

CM, CF electrophysiological and Ca^2+^ dynamics measurements; SB, ASM, ES biochemical measurements; CM, FL and AZ study design and manuscript writing; GMDF and AZ project and study coordination.

## References

1. Peichl P, Wichterle D, Pavlu L, Cihak R, Aldhoon B, Kautzner J. Complications of catheter ablation of ventricular tachycardia: a single-center experience. Circ Arrhythm Electrophysiol 2014;7:684–690.

2. Al-Khatib SM, Stevenson WG, Ackerman MJ, Bryant WJ, Callans DJ, Curtis AB, Deal BJ, Dickfeld T, Field ME, Fonarow GC, Gillis AM, Granger CB, Hammill SC, Hlatky MA, Joglar JA, Kay GN, Matlock DD, Myerburg RJ, Page RL. 2017 AHA/ACC/HRS Guideline for Management of Patients With Ventricular Arrhythmias and the Prevention of Sudden Cardiac Death: A Report of the American College of Cardiology/American Heart Association Task Force on Clinical Practice Guidelines and the Heart Rhythm Society. J Am Coll Cardiol 2018;72:e91–e220.

3. Zei P, Soltys S, Loo B, Norton L, Al-Ahmed A, Gardner E, Maguire P. First-in-man treatment of arrhythmia (ventricular tachycardia) using stereotactic radiosurgery. Heart Rhythm 2013;10.

4. Cuculich PS, Schill MR, Kashani R, Mutic S, Lang A, Cooper D, Faddis M, Gleva M, Noheria A, Smith TW, Hallahan D, Rudy Y, Robinson CG. Noninvasive Cardiac Radiation for Ablation of Ventricular Tachycardia. N Engl J Med 2017;377:2325–2336.

5. Robinson CG, Samson PP, Moore KMS, Hugo GD, Knutson N, Mutic S, Goddu SM, Lang A, Cooper DH, Faddis M, Noheria A, Smith TW, Woodard PK, Gropler RJ, Hallahan DE, Rudy Y, Cuculich PS. Phase I/II Trial of Electrophysiology-Guided Noninvasive Cardiac Radioablation for Ventricular Tachycardia. Circulation 2019;139:313–321.

6. Neuwirth R, Cvek J, Knybel L, Jiravsky O, Molenda L, Kodaj M, Fiala M, Peichl P, Feltl D, Januska J, Hecko J, Kautzner J. Stereotactic radiosurgery for ablation of ventricular tachycardia. Europace 2019;21:1088–1095.

7. Zhang DM, Navara R, Yin T, Szymanski J, Goldsztejn U, Kenkel C, Lang A, Mpoy C, Lipovsky CE, Qiao Y, Hicks S, Li G, Moore KMS, Bergom C, Rogers BE, Robinson CG, Cuculich PS, Schwarz JK, Rentschler SL. Cardiac radiotherapy induces electrical conduction reprogramming in the absence of transmural fibrosis. Nat Commun 2021;12:5558.

8. Zaza A, Rocchetti M. The late Na+ current--origin and pathophysiological relevance. Cardiovasc Drugs Ther 2013;27:61–68.

9. Stringer C, Pachitariu M. Cellpose3: one-click image restoration for improved cellular segmentation. Nat Methods 2025;22:592–599.

10. Berg S, Kutra D, Kroeger T, Straehle CN, Kausler BX, Haubold C, Schiegg M, Ales J, Beier T, Rudy M, Eren K, Cervantes JI, Xu B, Beuttenmueller F, Wolny A, Zhang C, Koethe U, Hamprecht FA, Kreshuk A. ilastik: interactive machine learning for (bio)image analysis. Nat Methods 2019;16:1226–1232.

11. Chamberland C, Barajas-Martinez H, Haufe V, Fecteau MH, Delabre JF, Burashnikov A, Antzelevitch C, Lesur O, Chraibi A, Sarret P, Dumaine R. Modulation of canine cardiac sodium current by Apelin. J Mol Cell Cardiol 2010;48:694–701.

12. Biet M, Morin N, Benrezzak O, Naimi F, Bellanger S, Baillargeon JP, Chouinard L, Gallo-Payet N, Carpentier AC, Dumaine R. Lasting alterations of the sodium current by short-term hyperlipidemia as a mechanism for initiation of cardiac remodeling. Am J Physiol Heart Circ Physiol 2014;306:H291–297.

13. Zhao Z, Fefelova N, Shanmugam M, Bishara P, Babu GJ, Xie LH. Angiotensin II induces afterdepolarizations via reactive oxygen species and calmodulin kinase II signaling. J Mol Cell Cardiol 2011;50:128–136.

14. Maltsev VA, Undrovinas A. Late sodium current in failing heart: friend or foe? Progress in Biophysics and Molecular Biology 2008;96:421–451.

15. Kleber AG, Rudy Y. Basic mechanisms of cardiac impulse propagation and associated arrhythmias. Physiol Rev 2004;84:431–488.

16. January CT, Riddle JM. Early afterdepolarizations: Mechanism of induction and block. A role for L-type Ca2+ current. Circulation Research 1989;64:977–990.

17. Zhao Z, Wen H, Fefelova N, Allen C, Baba A, Matsuda T, Xie LH. Revisiting the ionic mechanisms of early afterdepolarizations in cardiomyocytes: predominant by Ca waves or Ca currents? Am J Physiol Heart Circ Physiol 2012;302:H1636–H1644.

18. Zima AV, Blatter LA. Redox regulation of cardiac calcium channels and transporters. Cardiovasc Res 2006;71:310–321.

19. Kawamura K, Qi F, Kobayashi J. Potential relationship between the biological effects of low-dose irradiation and mitochondrial ROS production. J Radiat Res 2018;59:ii91–ii97.

20. Zeppenfeld K, Rademaker R, Al-Ahmad A, Carbucicchio C, De Chillou C, Cvek J, Ebert M, Ho G, Kautzner J, Lambiase P, Merino JL, Lloyd M, Misra S, Pruvot E, Sapp J, Schiappacasse L, Sramko M, Stevenson WG, Zei PC, Wichterle D, Chrispin J, Siklody CH, Neuwirth R, Pelargonio G, Reichlin T, Robinson C, Tondo C. Patient selection, ventricular tachycardia substrate delineation, and data transfer for stereotactic arrhythmia radioablation: a clinical consensus statement of the European Heart Rhythm Association of the European Society of Cardiology and the Heart Rhythm Society. Europace 2025;27.

