## Supplementary information for "Cellular consequences of non-ablative radiotherapy, a novel approach to ventricular tachycardias"

**SUPPLEMENT**

**DETAILED MATERIALS AND METHODS**

**Protein extraction and western blot analysis**

CMs were lysed in cell lysis buffer (Cell Signaling Technology, Danvers, MA, USA) supplemented with protease and phosphatase inhibitor cocktails (Sigma-Aldrich, Saint Louis, MO, USA). Total protein extracts were subjected to SDS-PAGE and transferred onto a nitrocellulose membrane (Bio-Rad, California, USA). The membranes were blocked for 1 h at room temperature in 5% non-fat dry milk in Wash Buffer (Tris Buffer Sulfate, 0.1% Tween-20) and then incubated O/N at 4 °C with the appropriate primary antibodies (reported in **Table S1**). The membranes were incubated with peroxidase-conjugated secondary antibodies (GE Healthcare, Chicago, IL, USA) for 1 h. Signals were visualized using the LiteUP Western Blot Chemiluminescent Substrate (EuroClone, Milan, Italy). Images were acquired with the ChemiDoc™ MP Imaging System (Bio-Rad, California, USA), and densitometric analysis of membranes was performed using the ImageJ software (National Institutes of Health, Bethesda, MD, USA). CMs CaMKII levels were normalized to GAPDH.

| **Protein** | **Clonality/Code** | **Source** | **Company** | **Diluition** |
| --- | --- | --- | --- | --- |
| phospho-CaMKII (T286) | Polyclonal, ab32678 | Rabbit | Abcam | WB: 1:1000 |
| CaMKII | Monoclonal, sc-5306 | Mouse | Santa Cruz | WB: 1:500 |
| GAPDH | Polyclonal, sc-25778 | Rabbit | Santa Cruz | WB: 1:1000 |

**Table S1.** Primary antibodies.

**SUPPLEMENTARY RESULTS**

|  | **CTRL** | **15 Gy** | **25 Gy** |
| --- | --- | --- | --- |
| Mean I_Nasus_ (pA/pF) | -0.45 ± 0.05 (n=18) | -0.79 ± 0.09 (n=17) | -1.53 ± 0.14 (n=16) |
| I_Nasus_/I_NaT_ (%) | 2.72 ± 0.21 (n=18) | 4.25 ± 0.59 (n=17) | 7.04 ± 1.20 (n=16) |
| I_NaT_ (pA/pF) | -18.43 ± 2.47 (n=18) | -20.69 ±1.82 (n=17) | -30.39 ± 4.53 (n=16) |
| E_rev_ | 45.62 ± 1.89 (n=18) | 46.68 ± 2.40 (n=17) | 43.71 ± 2.38 (n=16) |

**Table S2**. Summary of I_NaT_ and I_Nasus_ values.

|  | **CTRL** | **15 Gy** | **25 Gy** |
| --- | --- | --- | --- |
| APA (mV) | 101 ± 4.33 (n=16) | 109 ± 2.11 (n=17) | 113 ± 3.34 (n=19) |
| APD_90_ (ms) | 55.19 ± 7.27 (n=16) | 93.12 ± 13.71 (n=17) | 116 ± 16.79 (n=19) |
| +dV/dt_MAX_ (mV/ms) | 222 ± 27.09 (n=16) | 255 ± 15.34 (n=17) | 307 ± 27.68 (n=19) |
| E_diast_ (mV) | -77.01 ± 1.81 (n=16) | -80.39 ± 1.43 (n=17) | -74.93 ± 1.42 (n=19) |

**Table S3.** Summary of AP parameters.

|  | **CTRL** | **TTX** | **15 Gy** | **TTX** | **25 Gy** | **TTX** |
| --- | --- | --- | --- | --- | --- | --- |
| AP amplitude (mV) | 109 ± 5.81 (n=8) | 104 ± 5.86  (n=8) | 110 ± 1.94 (n=15) | 106 ± 2.15 (n=15) | 114 ± 3.74 (n=14) | 108 ± 4.82 (n=14) |
| APD_90_ (ms) | 36.22 ± 5.97  (n=8) | 33.93 ± 6.27 (n=8) | 98.23 ± 15.07 (n=15) | 87.16 ± 14.55 (n=15) | 119 ± 15.84 (n=14) | 89.55 ± 14.26 (n=14) |
| +/dt_MAX_ (mV/ms) | 280 ± 37.12 (n=8) | 238 ± 38.85 (n=8) | 264 ± 15.59  (n=15) | 237 ± 16.81 (n=15) | 316 ± 36.95 (n=14) | 251 ± 34.54 (n=14) |
| Diastolic potential (mV) | -79.76 ± 2.73  (n=8) | -76.62 ± 3.09 (n=8) | -80.46 ± 1.49 (n=15) | -80.22 ± 1.54 (n=15) | -75.69 ± 1.53  (n=14) | -76.61 ± 1.71 (n=14) |

**Table S4.** Summary of AP parameters before and after selective I_NaL_ blockade (1 M TTX).

|  | **CTRL** | **15 Gy** | **25 Gy** |
| --- | --- | --- | --- |
| CaD (F/F0) | 1.008 ± 0.004 (n=20) | 1.003 ± 0.004 (n=21) | 1.009 ± 0.004 (n=21) |
| CaT amplitude (F/F0) | 0.086 ± 0.011 (n=19) | 0.069 ± 0.007 (n=22) | 0.038 ± 0.005 (n=20) |
| tpeak (ms) | 86.75 ± 1.233 (n=20) | 90.02 ± 1.011 (n=21) | 95.04 ± 2.547 (n=21) |
| dCa/dt_MAX_ (F/ms) | 99.35 ± 4.033 (n=20) | 96.04 ± 2.616 (n=22) | 89.28 ± 1.949 (n=21) |
| τdecay (ms) | 523 ± 27.193 (n=20) | 551 ± 16.391 (n=22) | 625 ± 33.891 (n=21) |

**Table S5.** Summary of CaT parameters during steady-state stimulation at 1 Hz.

|  | **CTRL** | **15 Gy** | **25 Gy** |
| --- | --- | --- | --- |
| CaD (F/F0)_1Hz | 1.008 ± 0.004 (n=20) | 1.003 ± 0.004 (n=21) | 1.008 ± 0.004 (n=19) |
| CaD (F/F0)_5Hz | 1.049 ± 0.007 | 1.069 ± 0.008 | 1.078 ± 0.014 |
| CaT amplitude (F/F0)_1 Hz | 0.086 ± 0.011 (n=19) | 0.069 ± 0.007 (n=22) | 0.047 ± 0.010 (n=19) |
| CaT amplitude (F/F0)_5 Hz | 0.061 ± 0.006 | 0.052 ± 0.005 | 0.044 ± 0.009 |
| tpeak (ms)_1 Hz | 86.75 ± 1.233 (n=20) | 90.02 ± 1.011 (n=21) | 94.058 ± 2.575 (n=19) |
| tpeak (ms)_5 Hz | 59.34 ± 0.939 | 60.89 ± 1.708 | 66.82 ± 3.222 |
| dCa/dt_MAX_ (F/ms)_1 Hz | 99.35 ± 4.033 (n=20) | 96.04 ± 2.616 (n=22) | 89.55 ± 2.049 (n=19) |
| dCa/dt_MAX_ (F/ms)_5 Hz | 119 ± 4.116 | 122 ± 4.032 | 100 ± 5.759 |
| τdecay (ms)_1 Hz | 523 ± 27.193 (n=20) | 551 ± 16.391 (n=22) | 612 ± 35.683 (n=19) |
| τdecay (ms)_5 Hz | 384 ± 27.868 | 388 ± 32.047 | 359 ± 37.425 |

**Table S6.** Summary of CaT parameters during increasing stimulation frequency (from 1 Hz to 5 Hz).

|  | **CTRL** | **15 Gy** | **25 Gy** |
| --- | --- | --- | --- |
| pCaMKII vs CaMKII | 1 ± 0.106 (N=3) | 0.808 ± 0.006 (N=3) | 0.707 ± 0.018 (N=3) |

**Table S7.** Quantification by densitometric analysis of the pCaMKII/CaMKII ratio.

|  | **CTRL** | **25 Gy** |
| --- | --- | --- |
| ROS  (DCFDA fluorescence intensity, a.u.) | 175 ± 1.743 (n=16117) | 246 ± 2.469 (n=20708) |

**Table S8.** DCFDA fluorescence intensity.
